# The G-protein coupled receptor SRX-97 is required for concentration dependent sensing of Benzaldehyde in *Caenorhabditis elegans*

**DOI:** 10.1101/2020.01.04.894824

**Authors:** Nagesh Y. Kadam, Sukanta Behera, Sandeep Kumar, Anindya Ghosh-Roy, Kavita Babu

**Affiliations:** Department of Biological Sciences, Indian Institute of Science Education and Research (IISER) Mohali, Knowledge City, Sector 81, SAS Nagar, Manauli PO 140306, Punjab, India; Centre for Neuroscience, Indian Institute of Science, CV Raman Road, Bangalore 560012, Karnataka, India; National Brain Research Centre, Manesar, Nainwal Mode, Gurgaon 122051, Haryana, India

**Keywords:** SRX-97, ASH neuron, Benzaldehyde and C. *elegans*

## Abstract

The G-protein (heterotrimeric guanine nucleotide–binding protein)–coupled receptors in the olfactory system function to sense the surroundings and respond to various odorants. The genes encoding for the olfactory receptors in *C. elegans* are larger in number in comparison to those in mammals, suggesting complexity in the receptor– odorant relationships. Recent studies have shown that the same odorant in different concentration could act on multiple receptors in different neurons to induce attractive or repulsive responses. The ASH neuron is known to be responsible for responding to high concentrations of volatile odorants. Here we characterize a new GPCR, SRX-97. We found that the *srx-97* promoter shows expression specifically in the head ASH and tail PHB chemosensory neurons of *C. elegans*. Further, the SRX-97 protein localizes to the ciliary ends of the ASH neurons. Analysis of CRISPR/based deletion mutants of the *srx-97* gene suggest that this gene is involved in the recognition of high concentrations of benzaldehyde. This was further confirmed through rescue and neuronal ablation experiments. Our work gives insight into concentration dependent receptor function in the olfactory system and provides details of an additional molecule that could help the animal navigate its surroundings.

## Introduction

Animals sense a wide range of volatile and water-soluble chemicals through their olfactory system. The olfactory system is made up of several neurons that express different sets of seven-transmembrane G-protein coupled receptors (GPCRs). The odorant binds to the GPCRs, activating distinct intracellular signaling pathways and thus directing the animal’s response to different external cues (reviewed in (Erlandson et al., 2018; Katritch et al., 2013)).

*C. elegans* is a soil-dwelling animal that possesses a well-developed chemosensory system for their survival. They perceive their environment through various sensory neurons to find food sources, mates and to escape from dangerous conditions. In *C. elegans* 13 pairs of chemosensory neurons carry out majority of chemosensation, they express around 1300 functional chemosensory (cs) G-protein coupled receptors (GPCR) (Robertson and Thomas, 2006; Vidal et al., 2018). This diversity of csGPCRs allows the animal to discriminate between different odors. Thus, the specific expression of any GPCR or combined expression of different GPCRs on a specific neuron or different neurons can modulate the animal’s perception towards the same odorant.

The olfactory neurons that are involved in sensing a large number of attractive cues are AWA and AWC. These two pairs of neurons are involved in showing chemotaxis to various chemicals like diacetyl, isoamyl alcohol, pyrazine, benzaldehyde, and butanone (Bargmann et al., 1993; Colbert et al., 1997; Troemel et al., 1995). The avoidance behavior towards the repellents nonanone and 1-octanol is mediated through the sensory neurons AWB, ASH, and ADL (Chao et al., 2004). Besides this, many volatile chemicals detected by olfactory neurons could act as attractants at lower concentrations and repellents at higher concentrations (Yoshida et al., 2012). For example, at lower concentrations diacetyl is sensed by the GPCR ODR-10, in the AWA neuron acting as an attractant (Sengupta et al., 1996), while at higher concentration, it is sensed by the SRI-14 GPCR in the ASH neuron and acts as a repellent (Taniguchi et al., 2014). Additionally, ASH neurons are polymodal neurons involved in showing avoidance behavior towards different nociceptive signals like noxious chemicals, nose touch, hyperosmolarity, and volatile repellents (Bargmann, 2006; Hilliard et al., 2005). The ASH neurons convey information through different receptors. For example, touch has been shown to be detected by mechanically gated ion channels MEC-4 and MEC-10 while hyperosmolarity is detected by OSM-10 (Hart et al., 1999). The ASH neuron forms strong synaptic connections with the AVA command interneuron, which regulates backward locomotion of animals (Bhardwaj et al., 2019; Bhardwaj et al., 2018; Gray et al., 2005; Pokala et al., 2014; Zheng et al., 2012). Thus, activation of ASH neurons can affect backward locomotion or avoidance behavior in *C. elegans*.

ASH neurons are also reported to be involved in sensing undiluted or high concentrations of benzaldehyde (Aoki et al., 2011; Taniguchi et al., 2014; Troemel et al., 1995). Here, we show that SRX-97 a newly found csGPCR shows expression in the ASH neuron. By using the CRISPR/Cas9 method for genome editing, we made a deletion in the *srx-97* locus, generating a null mutation in *srx-97*. The *srx-97* mutants shows defect in chemotaxis behavior specifically towards high concentrations of benzaldehyde. Further, the mutant phenotype could be rescued by both endogenous and neuron-specific expression of the *srx-97* gene, suggesting concentration-dependent behavioral plasticity for odors in *C. elegans* through the SRX-97 GPCR.

## Materials and Methods

### *C. elegans* strains and maintenance

All *C. elegans* strains were maintained on nematode agar growth media (NGM) plates seeded with OP50 *Escherichia coli* at 20°C under standard conditions (Brenner, 1974). The *C. elegans*, N2 (Bristol strain) was used as the wild-type (WT) control and the mutant strains CX2205 *odr-3*(n2150) V, CX10 *osm-9*(ky10) IV, NL792 *gpc-1*(pk298) X, RB2464 *tax-2* (ok3403) I and VC3113 *tax-4*(ok3771) III used in this study were obtained from the *Caenorhabditis* Genetic Centre (CGC). Double mutants were made through standard genetic procedures and verified using PCR. The list of primers used for PCR verification in this study is tabulated in Table 1. The strains used in this study are listed in Table 2.

**Table 1:**
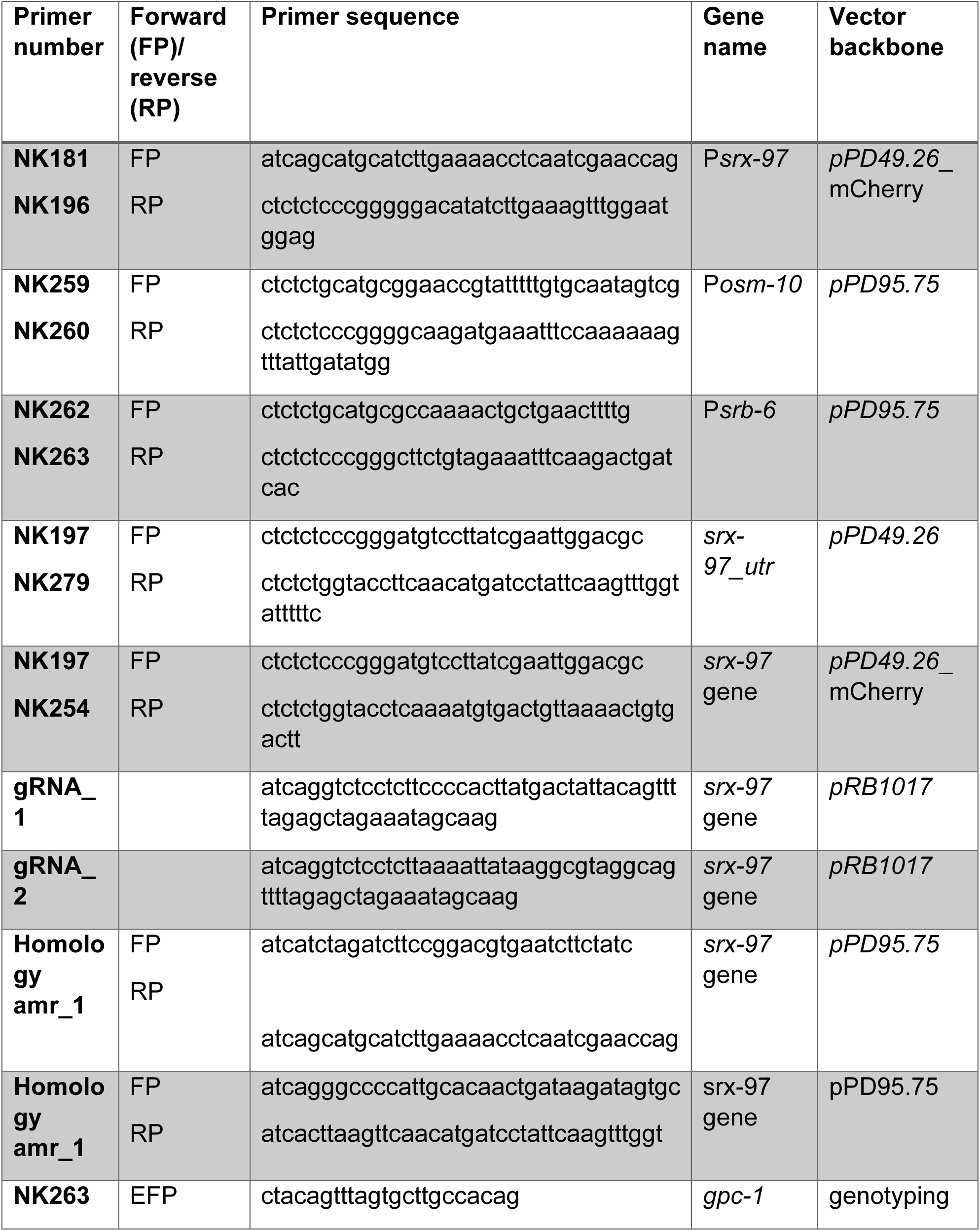
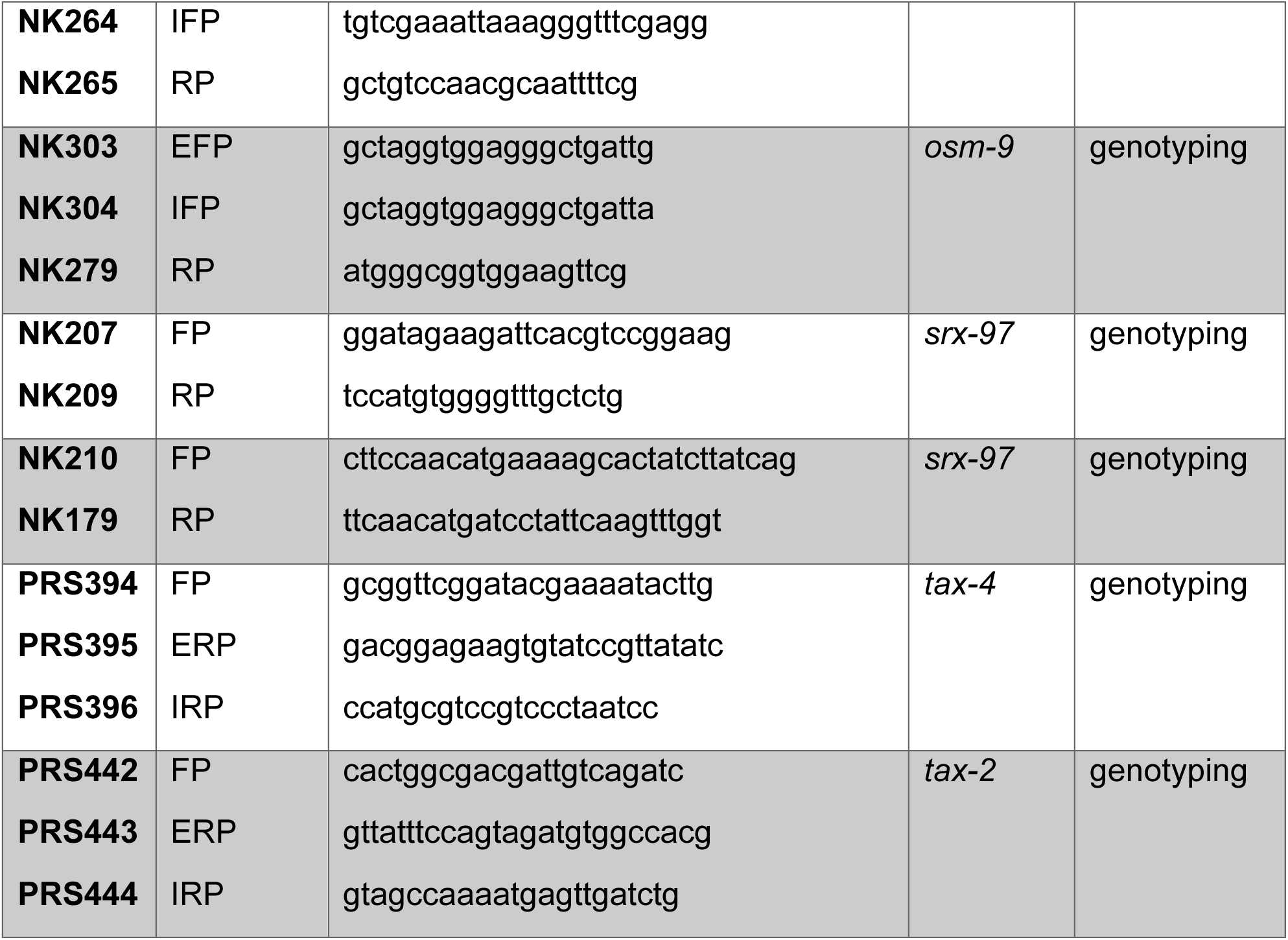
Primers used in this study.

**Table 2:** List of strain used in this study.

| Strain name | genotype | source |
| --- | --- | --- |
| **CX2205** | *odr-3* | CGC (3X outcrossed) |
| **CX10** | *osm-9* | CGC (3X outcrossed) |
| **RB2464** | *tax-2* | CGC (3X outcrossed) |
| **VC3113** | *tax-4* | CGC (3X outcrossed) |
| **NL792** | *gpc-1* | CGC (3X outcrossed) |
| **BAB404** | *srx-97* | This study (3X outcrossed) |
| **BAB482** | *srx-97;Psrx-97::SRX-97_utr (indEx462)* | This study |
| **BAB483** | *srx-97; Posm-10::SRX-97_utr (indEx466)* | This study |
| **BAB462** | *Psrx-97::SRX-97::mCherry (indEx462)* | This study |
| **BAB492** | *Psrx-97::mCherry (indEx460); Psrb-6::GFP (indEx465)* | This study |
| **BAB493** | *Psrx-97::mCherry (indEx460); Posm-10::GFP (indEx464)* | This study |
| **BAB473** | *srx-97;osm-9* | This study |
| **BAB487** | *srx-97;odr-3* | This study |
| **BAB488** | *srx-97;tax-2* | This study |
| **BAB489** | *srx-97;tax-4* | This study |
| **BAB490** | *srx-97;gpc-1* | This study |

### Rescue constructs and transgenes

All constructs for rescue of the *srx-97* phenotype were generated using standard cloning methods (Russell, 2001). The *pPD49.26* and *pPD95.75* vectors were used to clone the constructs. The primers used for cloning are indicated in Table 1. The transgenic lines were made using standard microinjection techniques as described previously (Mello and Fire, 1995; Mello et al., 1991). The *pCFJ90* and *pPD95.75* plasmids were used to amplify or clone mCherry and GFP respectively. The rescue constructs or promoter fusion constructs were injected at 20-30 ng/µl. P*myo-2*::mCherry (2 ng/µl) or P*unc-122*::GFP (25 ng/µl) were used as co-injection markers. The constructs used in this study are described in Table 3.

**Table 3:**
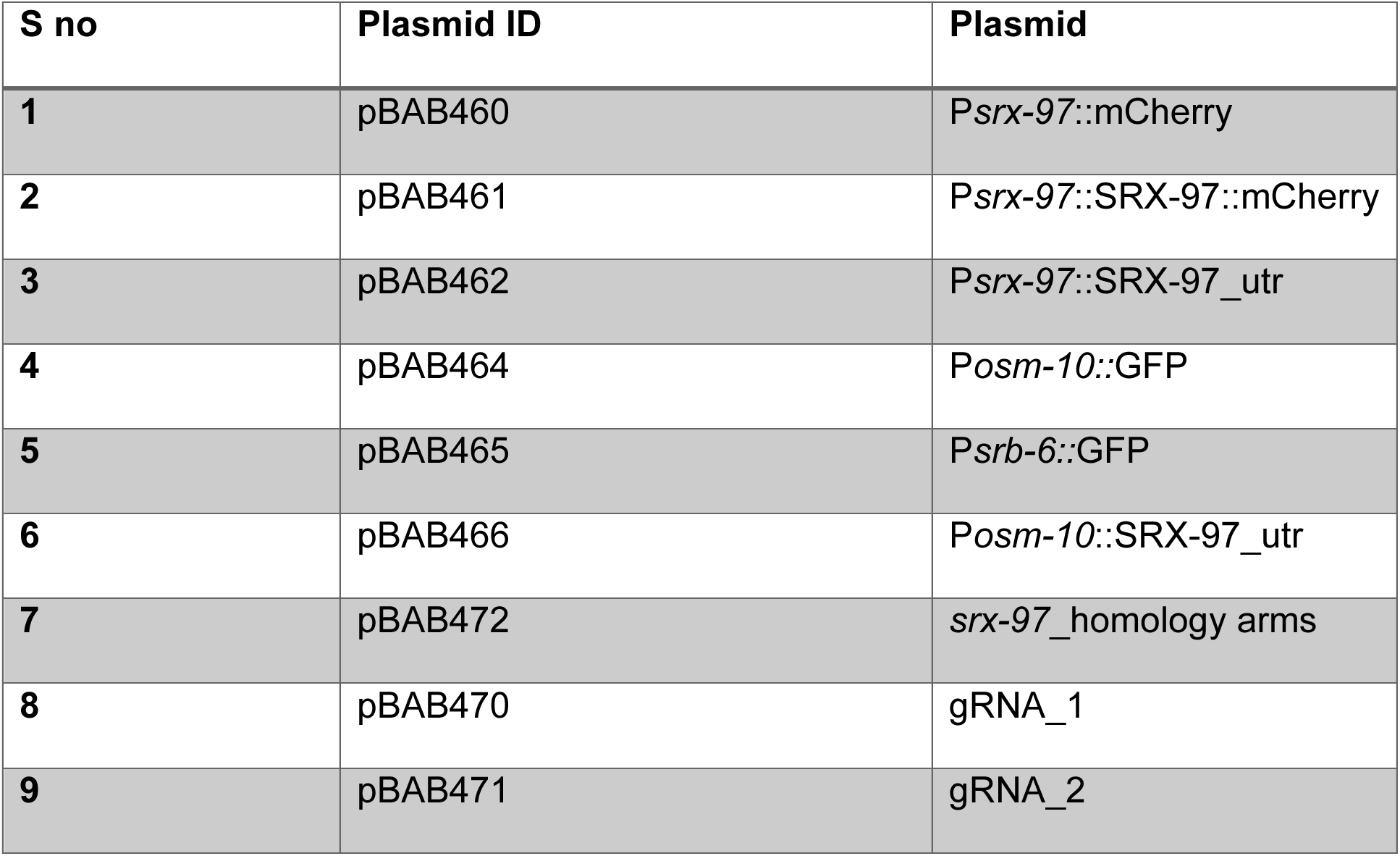
Plasmids used in this study.

| **S no** | **Plasmid ID** | **Plasmid** |
| --- | --- | --- |
| **1** | pBAB460 | *Psrx-97::mCherry* |
| **2** | pBAB461 | *Psrx-97::SRX-97::mCherry* |
| **3** | pBAB462 | *Psrx-97::SRX-97_utr* |
| **4** | pBAB464 | *Posm-10::GFP* |
| **5** | pBAB465 | *Psrb-6::GFP* |
| **6** | pBAB466 | *Posm-10::SRX-97_utr* |
| **7** | pBAB472 | *srx-97_homology arms* |
| **8** | pBAB470 | *gRNA_1* |
| **9** | pBAB471 | *gRNA_2* |

### Imaging experiments

Young adult animals were used for imaging. The animals were immobilized with 30 mg/ml 2, 3-butanedione monoxamine (BDM) on 2% agarose pads in M9 media. The promoter mCherry images for ASH and PHB were acquired on a Leica SP6 upright laser scanning confocal microscope using the 40X oil-immersion objective lens. Laser lines from He-Ne (594) with HyD detectors were used to image fluorescence in the head and tail regions. All other imaging experiments was done with a 40X or 63X 1.4 and 100X NA Plan APOCHROMAT objective using a Zeiss AxioCam MRm CCD camera on the Zeiss AxioImager Z2 microscope.

### Behavioral assays

#### Chemotaxis assay

The chemotaxis assay was done using young adult *C. elegans*. All chemotaxis assays were performed with standard 90 mm petriplates containing 15-18 ml of chemotaxis medium (Agar, MgSO_4_ (1M) CaCl_2_ (1M) and KPO_4_ (1M) at pH6.6). Wherever required, odorants were diluted in ethanol and reported as percent by volume. Modified 90 mm quadrant plate chemotaxis assays were performed as described previously (Bargmann et al., 1993; Margie et al., 2013). Briefly, 5 minutes (min) prior to the assay, 1 µl of 0.5 M Sodium azide was applied on four spots that were each 3 cm from the loading center. Sodium azide acts as an anesthetic agent to immobilize animals that reach the vicinity of the spot during the assay. Fifty to one hundred and fifty animals were placed at the center of the plate between the four spots, 2 µl of ethanol was placed at the two-control spots and 2 µl of the test odorant was placed at the two-test spots. After 90 min of chemotaxis, animals within each sector were counted, and the C.I. was calculated as the number of *C. elegans* in the two test sectors minus the number of animals in the two control sectors, divided by the total number of animals from each sector of the plate excluding the *C. elegans* that are not moving at the center of the plate (illustrated in Figure 3C). A positive Chemotaxis Index (C.I.) indicates an attraction to the chemical and a negative C.I. indicates an aversion to the chemical.

#### Assay to evaluate chemotaxis frequency

For analysis of the frequency or number of animals chemotaxing towards the source of benzaldehyde, a modified grid chemotaxis plate was used (Nuttley et al., 2001). The sodium azide was omitted so that animals could leave a spots after an initial approach. This grid consisted of four parallel lines drawn 1cm apart to divide the plate area into five sectors with the distance between the second and third lines being 2 cm (illustrated in Figure 4A). Two microliters of benzaldehyde were placed on one small sheet of parafilm and ethanol was placed on another as the control. The benzaldehyde and ethanol were placed at opposite ends of the plate (6 cm away). After a 60 min time interval, animals were immobilized by cooling the plates for 3 min at −30°C, and the plates were maintained at 4°C until counting. The number of animals in sectors a–d, with the test odorant being in a and d, were counted and a kinetic C.I. was calculated as (no. of animals in a + no. of animals in b) − (no. of animals in c + no. of animals in d) / (total number of *C. elegans* on the plate), yielding a C.I. range between +1.0 and −1.0. The *C. elegans* that crawl up the side of the plate were excluded from the analysis. The score for each plate of 50–150 animals is used as one data point.

#### Dry drop avoidance Assay

A drop of a solution containing the test chemicals (0.1SDS, quinine, CuSO4, glycerol, and pyrazine) dissolved in M13 (Tris, 30 mM; NaCl, 100 mM; KCl, 10 mM) buffer. The drop is delivered on the agar plate 0.5-1 mm anterior of the moving *C. elegans* (Hilliard et al., 2002). Once the animal encounters the dry drop of chemicals, the head amphid neurons sense the chemicals and the *C. elegans* shows repulsion or avoidance behavior. Normally WT C. *elegans* shows backward movement within 1 second (sec) while mutants like *odr-3* would show a significant delay in response. The delays in response to these different chemicals were calculated in the assay. Drops of M13 buffer were used as a control where animals as expected did not show any response. Glass capillaries (10 mm) pulled by hand on flames to reduce the diameter of the tip were used to deliver the drops.

### ASH neuron Ablation

ASH neuron ablation experiments were performed to test the benzaldehyde chemotaxis dependence on this neuron, which was tagged with P*srx-97*::mCherry. L2 staged animals were used for the ablation experiment, as loss of function is more effective in early stages (Avery and Horvitz, 1987, 1989; Bargmann and Horvitz, 1991). During ablation and imaging the animals were immobilized on 5% agarose pads with 0.1 μm-diameter polystyrene beads (00876-15; Polystyrene suspension). The Bruker Corporations ULTIMA setup was used to perform two-photon imaging and Ablation simultaneously (Basu et al., 2017). A 60X water immersion objective was used for ablation and imaging experiments, GFP and mCherry were imaged using 920 nm and 1040 nm lasers. A shot for 60 msec pulsed femtosecond IR laser [pulse width 80fs, irradiation pulse width: 50 msec, laser point spread function (PSF) 400 nm and Z axis PSF-1.5um] was used for all ablation experiments. Animals were then checked for proper ablation under a fluorescence microscope. These animals were allowed to grow and recover till they reach the young adult stage. Single animals were then transferred to unseeded plates and allowed to habituate for 1 min. Benzaldehyde (a a concentration of 10^-1^) was filled in the glass capillary having a small opening pore. The filled capillary was held just in front of the anterior region of the moving *C. elegans*. The lag time was calculated by considering the time taken by the animal to reverse half of its body length. Videos were recorded for 5 min at 10 frames/sec with 5-6 readings leaving a gap of about 1 min between each reading. The results were plotted using GraphPad Prism V6 and evaluated using *student t-test*, mean± SEM was plotted

### CRISPR/Cas9 mediated deletion of the *srx-97* gene

The <u>C</u>lustered <u>R</u>egularly Interspaced <u>S</u>hort <u>P</u>alindromic <u>R</u>epeats (CRISPR)/Cas9 system was used to create the *srx-97* deletion mutation as described previously (Dickinson and Goldstein, 2016). The two guide RNAs were designed (Hsu et al., 2013) and cloned separately into the *pRB1017* vector under the *CeU6* promoter. The Cas9 enzyme was expressed from the *pJW1259* vector under the *erf-3* promoter. The Selection Excision Cassette (SEC) containing plasmid *pDD287* was cloned along-with flanking loxP sites into the *pPD95.75* vector as described previously (Dahiya et al., 2019). The resulting plasmid was used to clone homology arms (500-600 bp) using restriction enzyme based cloning methods.

The plasmid mixture containing repair template (40 ng/µl), sgRNA_1 (10 ng/µl), sgRNA_2 (10 ng/µl), *pJW1259* (50 ng/µl), *pCFJ90* (2.5 ng/µl) and P*vha-6*::mCherry (15 ng/µl) was injected into 20-30 adult hermaphrodite animals (containing 4-5 eggs) that were kept at 20°C. Hygromycin was added after 60 hours (hr) of injection, directly on the NGM plate containing *C. elegans*. The hygromycin treated plates were left for 10 days at 20°C. Next 20-30 non-fluorescent rollers were singled out on normal seeded NGM plates. Once 100% roller progeny were observed on the plates these plates were kept at 34°C for 3-4 hr. Normally moving *C. elegans* were then picked and allowed to produce progeny. The genomic DNA was isolated from these progenies and the desired deletion was confirmed using PCR and sequencing techniques.

### Statistical analysis

All statistical analyses on behavioral assays were performed by using GraphPad Prism Version 6.0. The error bars represent SEM. Statistical significance was determined using a two-tailed unpaired Student’s *t-test* in figures 3B, 3D, 3E and 4C. For all other behavioral assays the data were compared using one-way ANOVA along with the Sadak’s *post hoc* test for multiple comparisons. Asterisks in the graphs indicate that the mean differences were statistically significant (*p* < 0.05). The level of significance were set as “*” *p* < 0.05; “**” *p* < 0.01; “***” *p* < 0.001.

## Results

### The P*srx-97*::mCherry transgene shows unique expression in the ASH and PHB chemosensory neurons

Chemosensory GPCRs are classified into nine different classes based on their sequence homology with the Rhodopsin class of molecules (Fredriksson et al., 2003; Lagerstrom and Schioth, 2008). The *C. elegans* genome has 1341 GPCR encoding genes, however the expression pattern of only 320 genes is known at a single-cell resolution (Robertson and Thomas, 2006; Taniguchi et al., 2014; Vidal et al., 2018). Reports suggest that GPCRs are also expressed in nonneuronal tissues like intestine and are involved in sensing internal cues (Vidal et al., 2018). Some GPCRs change their expression pattern once the animal encounters starvation or dauer like conditions (Vidal et al., 2018). Despite a large number of studies on the functional and spatial diversity of GPCRs, the expression pattern and function for a majority of the csGPCRs is still unknown (Robertson and Thomas, 2006; Taniguchi et al., 2014; Vidal et al., 2018).

We started this study to gain insight into the functioning of previously uncharacterized GPCRs and came across SRX-97. To determine the expression pattern of SRX-97, a region 2 kb upstream of the predicted translational start codon of the *srx-97* gene along with six base pairs of the first exonic region were used as a promoter to generate the P*srx-97::*mCherry reporter line. In these transgenic animals, mCherry expression was detected specifically in a single head amphid neuron and a tail phasmid neuron (Figure 1A-C). The amphid and phasmid neurons are involved in chemotaxis. No expression was detected in any other part of the body, suggesting that SRX-97 may specifically be involved in these neurons and is likely be involved in chemosensory signaling.

**Figure 1.**
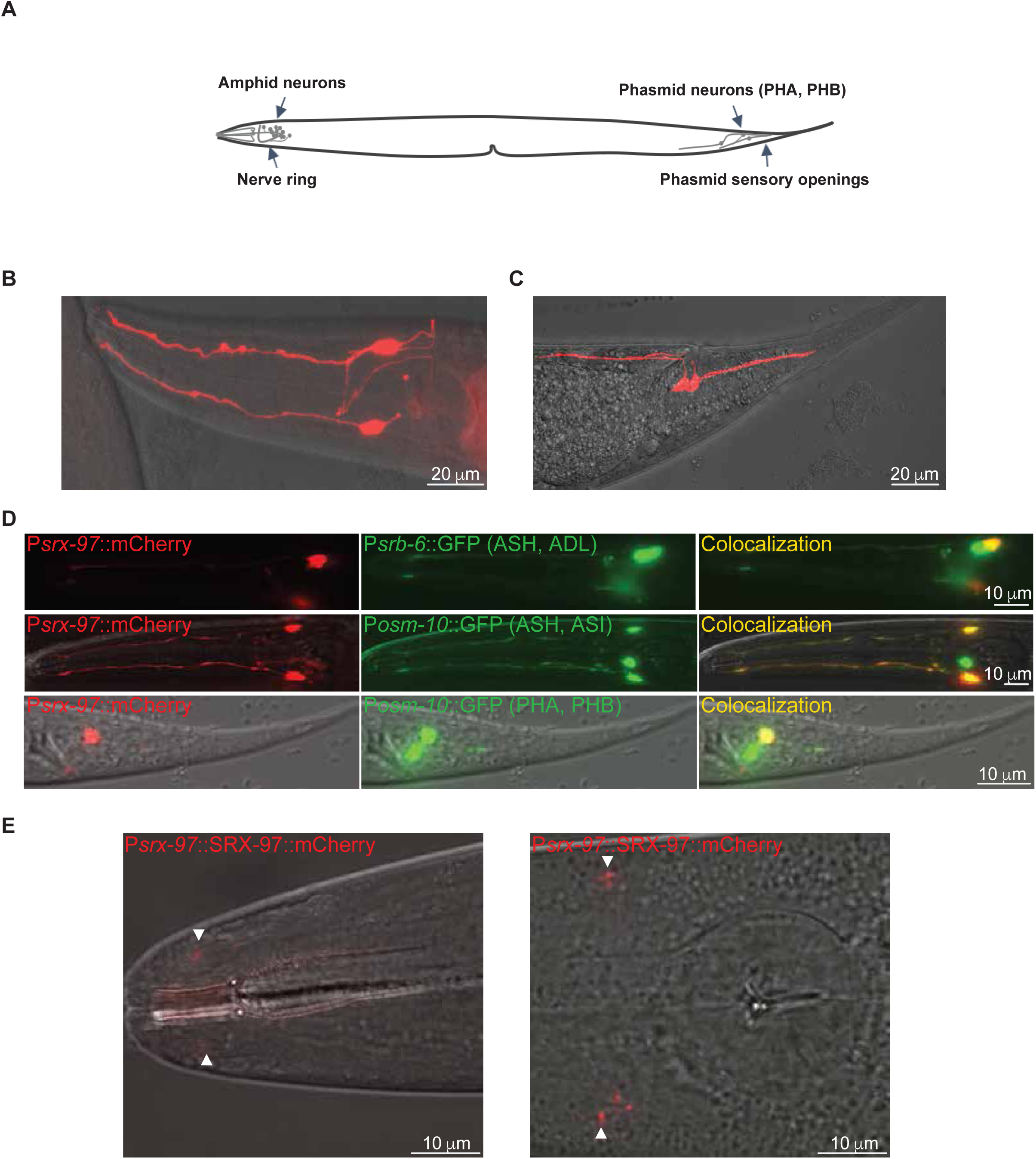
Expression of P*srx-97*::mCherry in ASH and PHB neuron: (A) A cartoon image showing the location of the amphid and phasmid neurons in *C. elegans.* (B) Shows expression of the P*srx-97*::mCherry transgenic construct in the amphid region and (C) The phasmid region of animal. (D) The expression of P*srb-6*::GFP and P*osm-10*::GFP in their respective (indicated on the figure) neurons and co-localization with P*srx-97*::mCherry in the amphid ASH and Phasmid PHB neuron. (E) The expression of SRX-97::mCherry in the cell body and it’s localization to the cilium tip of the ASH neuron.

Next, we began identifying the neurons expressing P*srx-97*:: mCherry based on cilium morphology, the cell body position in the head and tail regions and colocalization experiments. We first made the P*srb-6*::GFP transgenic line, which shows expression in the ASH and ADL neurons in the amphid region (Troemel et al., 1995). The P*srx-97::*mCherry line showed colocalization in a single neuron with the P*srb-6*::GFP line (Figure 1D, top panel). This suggested that P*srx-97*::mCherry could be expressed in either the ASH or the ADL neuron. To identify the P*srx-97*::mCherry expressing neuron, we made another transgenic line with P*osm-10*::GFP which shows expression in the amphid ASH and ASI neurons and the PHA/PHB neurons in phasmid region ((Figure 1D, middle panel) and (Hart et al., 1999)). The colocalization of P*srx-97*::mCherry in a single amphid neuron with both marker lines indicates that the *srx-97* promoter shows expression in the ASH neuron. In line with a recent report that suggests that 50% of GPCRs which express in ASH neurons also show expression in the PHB neuron (Vidal et al., 2018), we found that in the tail region, P*srx-97*::mCherry showed colocalization with the second phasmid neuron i.e., PHB neuron (Figure 1D, bottom panel).

We then went on to analyze the SRX-97 translational reporter and found that the P*srx-97*::SRX-97::mCherry transgenic line showed SRX-97 protein localization towards the cilium tip of the ASH neurons (Figure 1E), indicating that this protein may be involved in sensing environmental cues from the surroundings.

### CRISPR/Cas9 mediated deletion of *srx-97*

*C. elegans* have 13 pairs of chemosensory neurons in the anterior amphid and posterior phasmid regions, however, it can detect several different chemical cues ranging from volatile to water-soluble odorants through diverse GPCRs (Robertson and Thomas, 2006; Vidal et al., 2018). Seven percent of the *C. elegans* genome encodes for chemoreceptors. However, only around 900 chemoreceptor genes have been characterized functionally, many through RNAi experiments (Robertson and Thomas, 2006; Taniguchi et al., 2014; Vidal et al., 2018). Hence, only few mutant lines of GPCRs are available. Our expression studies showed that *srx-97* shows expression in ASH and PHB neurons. This expression pattern raised the possibility that it might function as a receptor for odorant/s. Since no mutant strain was available for this gene, we utilized the CRISPR/Cas9 based strategy to generate a deletion in the *srx-97* gene.

The SRX-97 GPCR is part of the SRX family of proteins that belong to the SRG superfamily that encodes around 320 genes (Robertson and Thomas, 2006; Vidal et al., 2018). The *srx-97* gene encodes a predicted protein of 317 amino acids (Figure 2A). Hydrophobicity analyses showed that the SRX-97 protein encodes for a seven-transmembrane domain protein, showing the characteristic topology of GPCRs (Figure 2A). By utilizing the gene editing CRISPR/Cas9 technique we made a complete deletion of the *srx-97* gene (from 61 bp of the first exon to the 3’ UTR region, deleting a 1834 bp sequence) (Figures 2 B and C).

**Figure 2.**
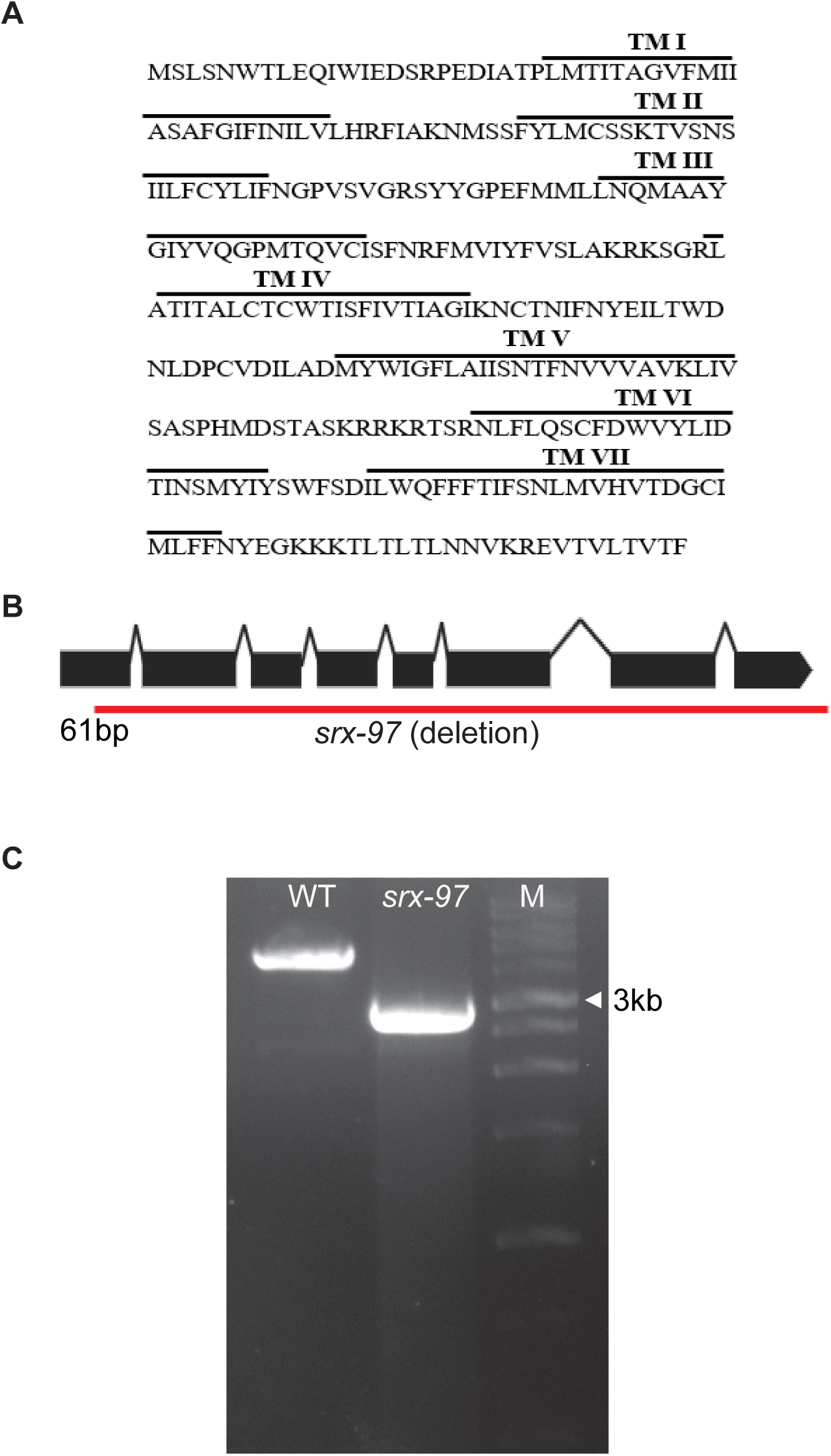
The SRX-97 transmembrane domain and CRISPR/Cas9 generated mutation of *srx-97*: (A) The amino acid sequence showing the predicted seven transmembrane domains of SRX-97. (B) The exonic structure of the *srx-97* gene with the red line showing the CRISPR/Cas9 deletion obtained. The deletion encompasses the gene from the 61^st^ base pair till the 1895^th^ base pair that is in the 3’ UTR of the gene. (C) Amplification of the chromosomal region showing the deletion of the *srx-97* gene (2730bp) using CRISPR/Cas9 compared to control wild-type (WT, 4566bp) *srx-97* gene. A 1kb DNA ladder was used in the line marked Marker (M).

### Loss of *srx-97* shows defects in chemotaxis towards Benzaldehyde

ASH is a polymodal neuron, it can respond to noxious, mechanical and osmotic stimuli (Colbert et al., 1997; Hilliard et al., 2005; Hilliard et al., 2004; Kaplan and Horvitz, 1993). To characterize the role of the SRX-97 GPCR in the ASH neuron, we checked the response of the *srx-97* mutant line towards several compounds like SDS, Cu^2+^, quinine, glycerol, pyrazine and acetic acid (Hilliard et al., 2005; Hilliard et al., 2002; Kaplan and Horvitz, 1993). The *srx-97* mutant animals did not show any significant defects in avoidance assays towards these compounds when compared to the control wild-type (WT) *C. elegans* (Figure 3A, B and Table 4).

**Figure 3.**
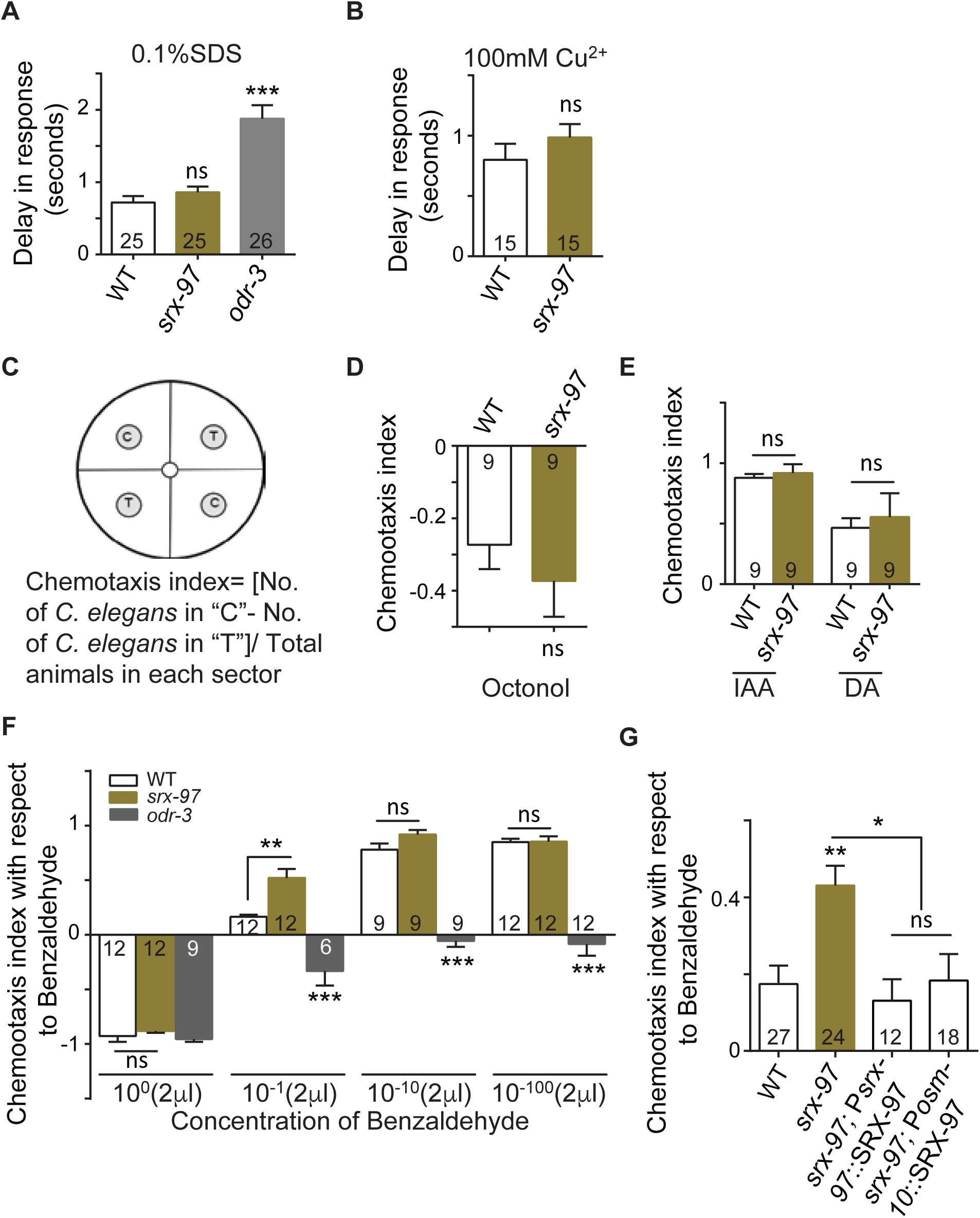
Behavior of *srx-97* mutant animals towards water soluble and volatile chemicals: (A) Graph showing the delay in avoidance towards a dry spot of 0.1% SDS and (B) towards the dry spot of 100mM CuSO_4_ in WT and *srx-97* mutant animals. The numbers at the base of graphs A and B indicate the number of animals tested for each genotype. (C) Schematic of a plate showing four quadrants, two opposite quadrants shows the control spots (termed C) and the test spots (termed T), 50-150 animals are added in the central spot and the chemotaxis index for volatile chemicals calculate by using the indicated formula. (D) Graph indicating the negative chemotaxis indices of WT and *srx-97* toward the repellent octanol. The assay was done in triplicates over multiple days. (E) Chemotaxis indices towards higher concentrations (10^-1^) of Diacetyl (DA) and Isoamyl alcohol (IAA). The assay was done in triplicates over multiple days. (F) Chemotaxis indices towards multiple concentrations of Benzaldehyde. The assay was triplicates in triplicates over multiple days. (G) Chemotaxis indices towards high concentrations of benzaldehyde in WT, *srx-97* and rescue strains of *srx-97*. The assay was done in triplicates over multiple days. The error bars represent SEM and statistical significance is represented as “ns” for not significant, “*” *p*<0.05, “**” *p*<0.01 and “***” *p*<0.001. The numbers at the base of each graph from D to G indicates the number of times the experiment was performed with 50-150 animals used in each trial.

**Table 4:** srx-97 mutant response towards different water soluble chemicals.

| Water soluble Chemicals | Genotype | Avoidance in second |
| --- | --- | --- |
| **Acetic acid**<br>**(0.1M)** | Wild type | 1.70±0.15 (n=21) |
|  | *srx-97* | 1.57±0.10 (n=20) |
| **Glycerol**<br>**(2M)** | Wild type | 1.30±0.8 (n=21) |
|  | *srx-97* | 1.20±0.6 (n=21) |
| **Dihydrocaffeic acid**<br>**(1M)** | Wild type | 0.60±0.01(n=20) |
|  | *srx-97* | 0.80±0.1 (n=21) |
| **Pyrazine**<br>**(10mM)** | Wild type | 1.0±0.40 (n=18) |
|  | *srx-97* | 1.20±0.30 (n=20) |
| **Quinine**<br>**(10mM)** | Wild type | 0.70±0.20 (n=15) |
|  | *srx-97* | 1.00±0.06 (n=21) |

The ASH neurons are also known to be involved in detecting volatile chemicals (Troemel et al., 1995). To analyze the role of SRX-97 in detecting volatile chemicals, we used a modified chemotaxis plate, having four quadrants, two opposite quadrants for test solutions (T) and two for control solutions (C) (illustrated in Figure 3 C). Both control and test spots were 3 cm away from the *C. elegans* loading center. Before the addition of control or test solution, we added sodium azide that paralyzes the animals once they reach the respective spots. Next, we calculated the chemotaxis index by measuring the number of worms in each quadrant with the formula shown in Figure 3C. Previous work has shown that in chemotaxis assays, the ASH neuron is involved in showing aversive behaviors towards the repellant 1-Octanol (Chao et al., 2004). Here again, we found no significant change in the chemotaxis index of the *srx-97* mutant line when compared to WT control animals (Figure 3D). Recent findings suggest that the ASH neuron is involved in sensing high concentrations of chemicals such as isoamyl alcohol (IAA) (Yoshida et al., 2012), and diacetyl (DA) (Taniguchi et al., 2014). In the chemotaxis assays, we used a range of concentrations of IAA and DA, testing for any defects in responses towards these chemicals. We found that the *srx-97* mutants did not show any significant defects in chemotaxis towards IAA and DA when compared to control *C. elegans* (Figure 3E and Table 5).

**Table 5:** ***srx-97* mutant response towards different concentration of volatile chemicals**

| Volatile chemicals | *srx-97*(CI) | WT(CI) | P value |
| --- | --- | --- | --- |
| **Diacetyl (10<sup>-10</sup>)** | 0.82 | 0.80 | 0.33 |
| **Diacetyl (10<sup>-100</sup>)** | 0.84 | 0.85 | 0.99 |
| Isoamyl alcohol ($10^{-10}$) | 0.83 | 0.84 | 0.33 |
| Isoamyl alcohol ($10^{-100}$) | 0.92 | 0.94 | 0.33 |

The ASH neurons are known to be involved in detecting benzaldehyde (Taniguchi et al., 2014; Troemel et al., 1995). A previously identified GPCR, DCAR-1 has homology with the SRX family of proteins. Further, *dcar-1* mutants show defective chemotaxis towards undiluted benzaldehyde (Aoki et al., 2011). In our chemotaxis assay, the *srx-97* mutant animals showed significantly more attraction towards a high concentration of benzaldehyde (10^-1^) when compared to WT controls (Figure 3F). At lower concentrations (10^-10^ and 10^-100^) of benzaldehyde and undiluted benzaldehyde, there was no significant difference between *srx-97* and WT animals (Figure 3F). It has been previously reported that the ASH neuron is involved in responding to high concentrations of benzaldehyde (0.1% v/v), while medium or lower concentrations (0.005-0.0001%) of benzaldehyde are sensed by the AWC and AWA neurons (Leinwand et al., 2015). Since SRX-97 is expressed in the ASH neuron, it could be involved in sensing a very specific concentration range of Benzaldehyde. In order to confirm the *srx-97* mutant phenotype, we went on to rescue the defects seen in the *srx-97* mutant line. We found that the defects in chemotaxis towards benzaldehyde seen in the *srx-97* animals could be rescued by expressing SRX-97 under its native promoter as well as with the *osm-10* promoter that shows expression in the ASH and ASI neurons (Figure 3G). These data suggest that the csGPCR SRX-97 is responsible for sensing the high concentrations of benzaldehyde.

### Ablation of ASH causes defects in Benzaldehyde sensing

We next analyzed the chemotaxis frequency of *srx-97* mutants towards high concentrations of benzaldehyde (Nuttley et al., 2001). Here we added the benzaldehyde (10^-1^) on a small sheet (0.5-1 cm diameter) of parafilm so it would not be soaked in the media. We also excluded the addition of sodium azide on the control and test spots so as to allow the *C. elegans* to move freely towards the control or test spot (Illustrated in Figure 4A). After a 60 min incubation period, the number of animals were counted sector-wise and the chemotaxis frequency was calculated by the formula indicated in Figure 4A. Again the *srx-97* mutants showed a significant increase in their attraction towards benzaldehyde (Figure 4B). This defect was rescued by expressing SRX-97 under its endogenous promoter, again suggesting the SRX-97 is responsible for sensing high concentrations of benzaldehyde.

**Figure 4.**
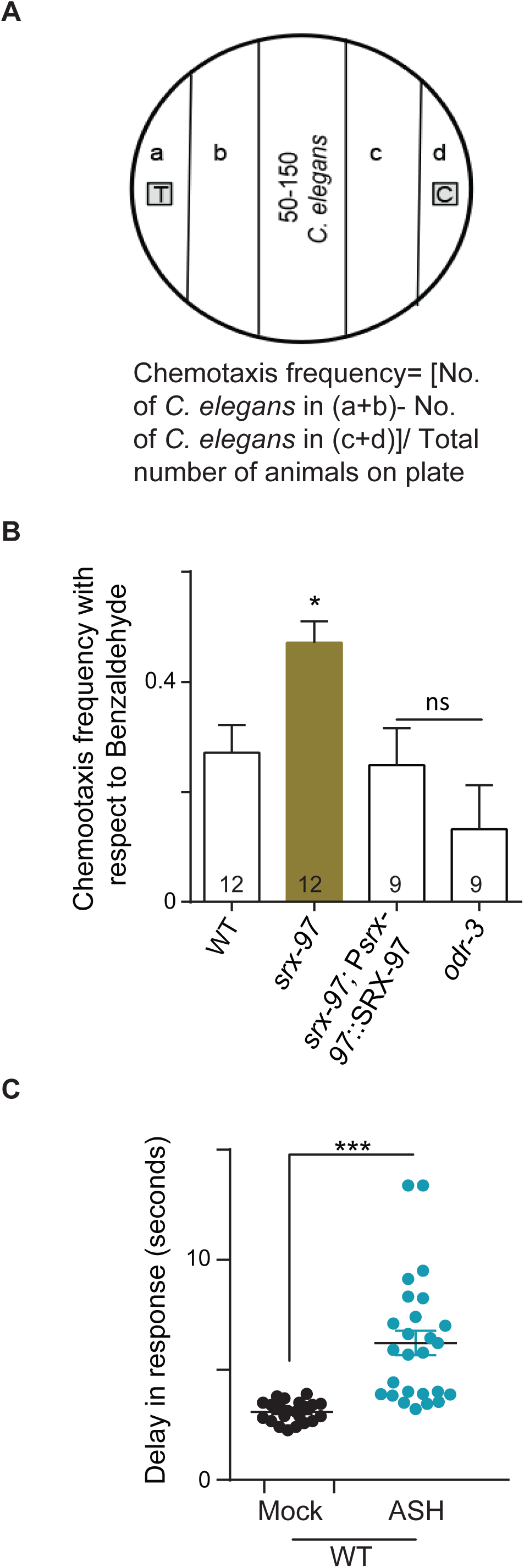
ASH neuron ablated animals’ shows defects towards chemosensation to benzaldehyde: (A) An illustration of the design of the plates used for analyzing the chemotaxis frequency of *C. elegans* along with the formula used for this calculation. Each sector (a,b,c and d) is 1 cm in width. (B) Graph of chemotaxis frequencies of WT, *srx-97*, the *srx-97* rescue line and a control *odr-3* mutant line to a high concentration of benzaldehyde. The assays were performed in triplicate over multiple days and the numbers at the base of each graph indicates the number of times the experiment was performed with each genotype. (C) Graph plotting the delay in response of WT animals that have undergone mock ablation or ASH ablation towards high benzaldehyde concentration. Each dot indicates a response from a single animal. 22 mock ablated animals and 26 ASH ablated animals were analysed for this experiment. The error bars represent SEM and statistical significance is represented as “ns” for not significant, “*” *p*<0.05 and “***” *p*<0.001.

In order to gain further evidence that SRX-97 was indeed acting in ASH to sense high concentrations of benzaldehyde, we ablated the ASH neuron in WT animals. We then went on to test the delay in response towards benzaldehyde (10^-1^) in mock ablated and ASH ablated *C. elegans*. Our data showed that the ASH neuron ablated WT animals showed significant delay in their response to benzaldehyde when compared to the mock-ablated animals (Figure 4C).

### Defects in sensory signaling appear to function downstream of *srx-97*

The ASH neurons expresses multiple GPCR associated sensory molecules that are reported to be required for signal transduction (Hilliard et al., 2005; Hilliard et al., 2004; Roayaie et al., 1998). Among these the G protein subunit ODR-3 is a Gα protein which is primarily required for sensory signal transduction and is involved in response towards osmotic strength, high salt concentration, nose touch and volatile chemicals (Hilliard et al., 2005; Hilliard et al., 2004; Roayaie et al., 1998; Zhang et al., 2016). Mutants in *odr-3* have been reported to show attraction towards low concentrations of benzaldehyde (1:200) (Roayaie et al., 1998). Likewise mutants in *gpc-1* that encodes the γ subunit of GPCRs show positive adaptive olfactory responses towards benzaldehyde (Jansen et al., 2002; Yamada et al., 2009). In our assay, we found that mutants of both these GPCRs associated sensory molecules and also the double mutants with *srx-97* showed negative chemotaxis indices towards high concentration of benzaldehyde similar to what was seen with the *odr-3* and *gpc-1* single mutant animals (Figure 5). OSM-9 is a member of the vanilloid subfamily of Transient Receptor Potential (TRP) channel proteins that regulates avoidance behavior to osmotic strength, nose touch and undiluted benzaldehyde in the ASH neuron (Colbert et al., 1997; Murayama and Maruyama, 2013; Zou et al., 2017). We found that both the single *osm-9* mutant and double mutants of *osm-9* and *srx-97* showed wild type like phenotype to high concentration of benzaldehyde (Figure 5). Finally, we tested the cyclic nucleotide-gated channel proteins TAX-2 and TAX-4. These proteins are responsible for detection of volatile chemicals like benzaldehyde by AWC and other amphid neurons although the source of activating cGMP is still unknown (Coburn and Bargmann, 1996; Komatsu et al., 1996; Zagotta and Siegelbaum, 1996). These mutants were also used on quadrant plate chemotaxis assays and showed a negative chemotaxis index toward high concentration of benzaldehyde (Figure 5). Again, the double mutant of *tax-2* and *tax-4* with *srx-97* behaved like the single *tax-2* and *tax-4* mutants. Suppression of the *srx-97* mutant phenotype by these downstream molecules suggest that either SRX-97 GPCR acts redundantly to sense the high concentrations of benzaldehyde by activating pathways different from the ones tested or there is a possibility that these downstream molecules function to detect both high and low concentrations of benzaldehyde and SRX-97 functions through the canonical G-protein pathway to elicit responses to high concentrations of benzaldehyde.

**Figure 5.**
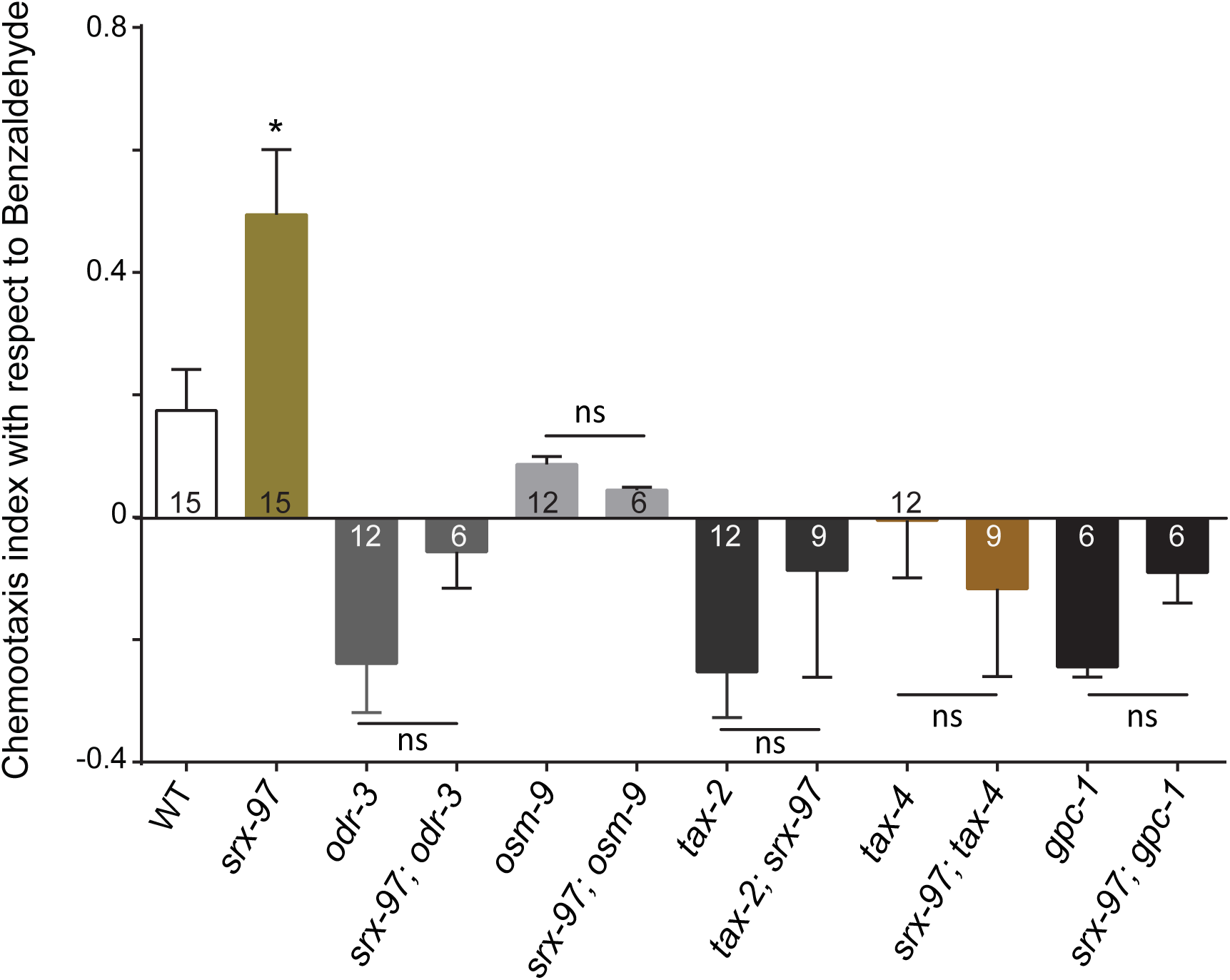
The *srx-97* mutant phenotype is suppressed by other signaling mutants that appear to function downstream of SRX-97: Chemotaxis indices with respect to high concentration of benzaldehyde in WT, *srx-97, odr-3, osm-9, tax-2, tax-4* and *gpc-1* mutants along with analysis of each mutant in the srx-97 background. The assays were performed in triplicate over multiple days and the numbers at the base of each graph indicates the number of times the experiment was performed with each genotype.The error bars represent SEM and statistical significance is represented as “ns” for not significant and “*” *p*<0.05.

## Discussion

In this study we have characterized the expression and function of the GPCR, SRX-97. From our expression studies it is clear that SRX-97 shows expression in the ASH and PHB neurons. Further the chemotaxis experiments reveal that the GPCR SRX-97 senses high concentrations of benzaldehyde. Our data indicate that in comparison with wild-type (WT) animals, *srx-97* null mutant *C. elegans* show increased attraction towards high concentrations of benzaldehyde (10^-1^). We also show that SRX-97::mCherry driven by its native promoter shows localization towards the cilium tip of the ASH neuron. Since the cilia are the compartment where signal sensation and transduction occurs, the localization of SRX-97 at the cilium tips suggests its role in sensory perception or transduction of sensory signal/s. These results further suggest that SRX-97 expressed in the ASH neuron is responsible for detecting benzaldehyde from its surroundings.

Other than SRX-97, ASH neurons also express different sets of GPCRs, which sense benzaldehyde (Aoki et al., 2011; Taniguchi et al., 2014; Vidal et al., 2018). For example, DCAR-1 is expressed in ASH neurons and is involved in sensing undiluted benzaldehyde (Aoki et al., 2011). Since the *srx-97* mutant animals show reduced but not completely abolished response towards high concentrations (10^-1^) of benzaldehyde, it is possible that SRX-97 may act as a constituent of a receptor complex on the ASH neuron that may specifically detect benzaldehyde at a high or undiluted concentrations but not at low concentrations. Low concentration of benzaldehyde on the other hand is sensed by the AWC neuron (Bargmann et al., 1993; Leinwand et al., 2015). The wild-type like chemotaxis response of *srx-97* mutants towards undiluted and low concentration of benzaldehyde may suggest that animals sense their surrounding by activating different receptors using the corresponding neurons in a concentration dependent manner and this in turn leads to appropriate behavioral responses.

GPCRs signal through the heteromeric G-proteins signaling cascades and transduce signals from the environment through intracellular mediators that play an important role in triggering behavior. The ASH and other amphid neurons express the Gα protein ODR-3 as well as OSM-9, a TRPV protein that are involved in sensation of various stimuli including olfaction (Bargmann et al., 1993; Hilliard et al., 2004; Roayaie et al., 1998; Troemel et al., 1997). The amphid AWC neuron acts as a primary olfactory neuron involved in sensing low concentrations of benzaldehyde (Leinwand et al., 2015), while the ASH neuron is required for sensing undiluted benzaldehyde (Colbert et al., 1997; Tobin et al., 2002; Troemel et al., 1995). Our study assaying chemotaxis in C. elegans at 90 minutes found that *srx-97* mutants showed chemotaxis defects towards high concentrations of benzaldehyde, in these assays the animals were placed 3 cm away from the source of benzaldehyde and hence not in short range of the source (Troemel et al., 1995). Previous work has shown that the distance or diffusion gradient of a test chemical may activate primary sensory neurons like AWC and AWB (Leinwand et al., 2015; Taniguchi et al., 2014; Yoshida et al., 2012). Our work also indicates that defects in the downstream signaling molecules in these neurons could affect the repulsion of the animals from the source.

Our results further suggest that there could be alternative pathways for signal transduction in ASH neurons through GPCRs like SRX-97. To our knowledge, not a single downstream signaling molecule has been identified the loss of which shows attraction to undiluted or high concentration of benzaldehyde through the ASH neuron in the chemotaxis assay. The *C. elegans* genome encodes for the 21 Gα, 2 Gβ and 2 Gγ genes (Cuppen et al., 2003; Jansen et al., 1999). Out of these 11 Gα proteins are known to express in ASH neurons (Bastiani and Mendel, 2006). Our results hypothesize that the ASH neuron is involved in aversion to undiluted or high concentration of benzaldehyde through multiple or redundant chemosensory pathways involved in the signaling through GPCRs like SRX-97.

In conclusion, our results bring out the possibility that SRX-97 is a key mediator in chemotaxis towards high concentrations of benzaldehyde in the chemosensory system of *C. elegans*. However, the downstream signaling components still needs to be deciphered, which can help in providing a better overview of the SRX-97 dependent pathway. Further these investigations may offer insights into the nature of signal transduction in ASH neurons and its physiological role in concentration dependent avoidance responses.

## Acknowledgments

The authors are especially grateful to Yogesh Dahiya for help with making the *srx-97* deletion strain. A number of strains were provided by CGC, which is funded by NIH Office of Research Infrastructure Programs (P40 OD010440). The authors thank Ankit Negi for routine help and the IISER Mohali Confocal facility for use of the confocal microscope. AG-R thanks Harjot Kaur for the help with 2-Photon microscopy.

NYK thanks Council of Scientific and Industrial Research (CSIR)-University Grants Commision (UGC) for a graduate fellowship and was also funded by a Department of Science and Technology (DST)-Science and Engineering Research Board (SERB) grant to KB. SB was funded by a KVPY fellowship for undergraduate students. KB was an Intermediate Fellow of the India Alliance (IA) and thanks the Alliance for funding support. KB also thanks DST-SERB and DBT for funding support.

## Author Contributions

NYK designed, performed, analyzed all the experiments and wrote the manuscript. SB helped with designing and performing experiments. SK and AG-R helped with performing the ablation experiments and editing the manuscript. KB supervised the experiments, helped with experimental design and data interpretation and edited the manuscript.

The authors declare no conflict of interest.

